# High-density Neural Recordings from Feline Sacral Dorsal Root Ganglia with Thin-film Array

**DOI:** 10.1101/2020.07.14.199653

**Authors:** Zachariah J. Sperry, Kyounghwan Na, James Jun, Lauren R. Madden, Alec Socha, Eusik Yoon, John P. Seymour, Tim M. Bruns

## Abstract

Objective: Dorsal root ganglia (DRG) are promising sites for recording sensory activity. Current technologies for DRG recording are stiff and typically do not have sufficient site density for high-fidelity neural data techniques. Approach: In acute experiments, we demonstrate single-unit neural recordings in sacral DRG of anesthetized felines using a 4.5 μm-thick, high-density flexible polyimide microelectrode array with 60 sites and 30-40 μm site spacing. We delivered arrays into DRG with ultrananocrystalline diamond shuttles designed for high stiffness affording a smaller footprint. We recorded neural activity during sensory activation, including cutaneous brushing and bladder filling, as well as during electrical stimulation of the pudendal nerve and anal sphincter. We used specialized neural signal analysis software to sort densely packed neural signals. Main results: We successfully delivered arrays in five of six experiments and recorded single-unit sensory activity in four experiments. The median neural signal amplitude was 55 μV peak-to-peak and the maximum unique units recorded at one array position was 260, with 157 driven by sensory or electrical stimulation. In one experiment, we used the neural analysis software to track eight sorted single units as the array was retracted ~500 μm. Significance: This study is the first demonstration of ultrathin, flexible, high-density electronics delivered into DRG, with capabilities for recording and tracking sensory information that are a significant improvement over conventional DRG interfaces.

## Introduction

Dorsal root ganglia (DRG) are neural structures with tremendous potential as bioelectrical interface sites, but current technologies available to access, map, and utilize the dense sensory information they contain are limited. As peripheral nerves enter the central nervous system, sensory neurons first coalesce at each spinal level into bilateral dorsal spinal nerves. These nerves, or dorsal roots, each contain a single ganglion, or DRG, which in turn contain the unmyelinated cell bodies of all sensory neurons entering that spinal level. When conducting an action potential, these cell bodies generate a relatively large extracellular potential detectable at single-unit fidelity by nearby recording electrodes [1]. The sensory information that can be decoded from these signals can be used as feedback to control, for example, neural stimulation for bladder control or walking [2]–[7].

However, much remains unknown about the intrinsic anatomy of DRG. Previous studies have presented some evidence of functional organization within individual DRG [8], [9], but the overall structure-function relationship still remains unclear. In comparison, functional organization relationships in the brain and spinal cord have been well-characterized, possibly allowing for the development of selectively targeted neural interfaces for particular applications. Presently DRG can be targeted to choose a ganglion at a particular spinal level, such as sacral DRG for bladder-related applications or lumbar DRG for lower limb neuroprostheses. Within-DRG interfacing for selective access to individual peripheral nerve pathways generally depends on the random nature of inserted microelectrodes being located near axons of interest, however. Tools to study the organization of DRG *in vivo* could lead to more selective targeting within these structures.

The current standard for *in vivo* recording of DRG neurons is the Utah array, a commercially available, silicon-based, penetrating microelectrode array. Previous studies have successfully demonstrated the capability of Utah arrays to record a variety of sensory neurons in the DRG, including populations related to urinary tract function, joint flexion, and skin sensation [2]–[7]. However, the mechanical mismatch between silicon and neural tissue causes tissue damage and scarring *in vivo* [10]. Floating microelectrode arrays (FMAs), which allow for custom shank lengths and tip impedances, have been used for DRG recording and stimulation with a minimum 250 μm site spacing [11]–[14], and also have scarring around electrode shanks for chronic *in vivo* implants [15]. DRG recordings have also been reported with a single-shank silicon probe (“Michigan probe”), with 50 μm electrode-site spacing [16]. The stiff nature of these probes is likely to cause a chronic tissue-scarring and immunological response, as has been reported in the brain and peripheral nerves including DRG [10], [15], [17], [18].

Another known issue with neural probes is “electrode drift,” the tendency of the electrode site to shift relative to the neuron of interest [19]. Because Utah arrays and FMAs have only a single recording site within any given neuron’s recordable range, drift due to scarring or micromotion makes them susceptible to loss of signals of interest or changes to the waveform that cause challenges during spike sorting [19]. A number of algorithmic methods are available to help mitigate the issue of waveform shape change [20]–[22]. Higher density probes may be preferable to reduce signal loss and simplify corrections by oversampling single neural units on multiple channels, and attention is increasingly focused on these high-density arrays as software becomes available to efficiently handle the quantity of data generated in single recordings [19], [23].

Based on these constraints, we believe that a flexible and high-density electrode array would be the preferred interface for mapping within DRG. One way to achieve this is with planar electrode arrays with a thin-film polymer substrate, first described for high-fidelity neural recording in the brain by Rousche et al. (2001) [24]. Thin-film polymer devices with thickness around 20 μm have been demonstrated to produce significantly less neuronal loss and glial response than larger devices on the order of 50 μm thickness [25]–[27]. We previously reported the use of a high-density non-penetrating polyimide array for single-unit neural recording from the surface of lumbosacral DRG [28], and studies have used other technologies to record from the DRG surface, [29]–[31] but biophysical limitations suggest that no units would be recorded below about 200 μm below the surface. Though anatomical analysis suggests that the highest density of somata reside in this outer dorsal region of the DRG [32], [33], selective mapping or microstimulation requires a technology interfacing with the interior of DRG.

In this study, we demonstrate high-density recording and mapping applications in sacral feline DRG using a flexible polymer array recently developed by Na et al. at the University of Michigan [34]. In this previous report, emphasis was placed on the specialized array delivery device, while in this report we will focus on DRG recordings. Together, these studies represent the first demonstration of flexible array recordings in DRG. The array was similar in design to the one reported in Sperry et al. (2017) [28], but was delivered into the DRG with a novel structurally-stiffened diamond shuttle. Sacral DRG were targeted because of their potential use as interface sites for bladder neuroprosthetic devices, though the technology could be directly transferred to other spinal levels or neural interface sites. We successfully delivered arrays in 5/6 acute experiments and recorded high-density sensory neural activity in 4 of these experiments. We used this high-density information to efficiently sort the neural signals and to track individual neurons as the array was moved through the DRG to simulate the extremes of chronic recording conditions.

## Methods

### Microelectrode Array

The primary purpose of this study was to explore the recording and mapping capabilities of a high-density microelectrode array in feline sacral DRG. Arrays were fabricated in the Lurie Nanofabrication facility utilizing the same process described for the ganglionic surface electrode arrays in Sperry et al. 2018 [28], but with modifications in the overall design. In brief, platinum electrode sites were patterned and connected with gold/platinum traces sandwiched in the middle of a 4.5 μm thick flexible polyimide substrate. In this study, each of the 60 sites were approximately square, with an area of 400 μm^2^, and arranged in 2 off-set columns. The pitch between electrodes was 40 μm. The active portion of the array was 1160 μm long and tapered from 80 μm wide to 55-μm wide for most of the length of the shank. Figure 1(a) shows a schematic of the array. The high-density array layout allowed for oversampling of units across electrode sites for enhanced sorting capabilities and tracking of units while the array was retracted.

**Figure 1:**
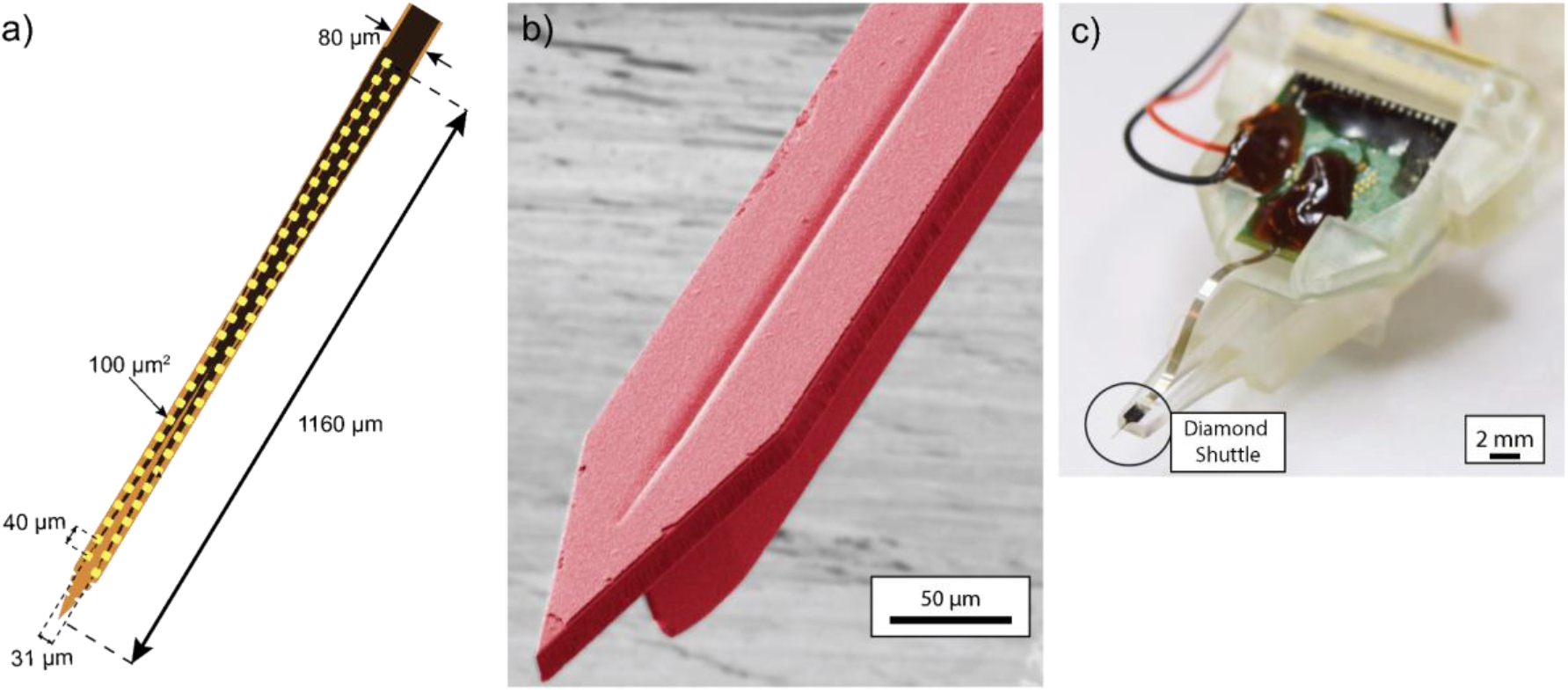
Flexible intraneural DRG array. (a) Tip of flexible high-density array showing locations of 60 electrode sites (yellow) and dimensions. (b) Diamond shuttle imaged with electron microscope, false color for visibility. (c) Insertion jig, with location of diamond shuttle highlighted in a circle.

Each array was bonded to a custom printed circuit board (PCB) for interfacing with the neural recording system. To reduce site impedance prior to recording, array channels were coated with poly(3,4-ethylenedioxythiophene) polystyrene sulfonate (PEDOT:pSS) as described in Patel et al., with the current adjusted for the electrode site area [35]. To verify all deposition and coating steps, impedance measurements were taken with a PGSTAT12 Autolab (EcoChemie, Utrecht, Netherlands), controlled by vendor supplied NOVA software. Measurements were obtained by applying a 1 kHz 10 mV_rms_ signal. Custom MATLAB (Mathworks, Natick, MA) scripts were used to determine frequency-specific impedance values. The PCB board was placed in a custom 3D-printed jacket and mounted to a 3D-printed insertion jig (Form2 3D printer, Formlabs, Somerville, MA) (Figure 1(c)).

For delivery into DRG, the flexible array was temporarily adhered to an ultrananocrystalline diamond (UNCD) shuttle with water-soluble polyethylene glycol (PEG; 12,000 MW) or ultraviolet-cured cyanoacrylate glue. The shuttle was fabricated with a stiffened T-profile by UNCD deposition over a trench which was etched away to form the final shape (fabrication details and characterization in Na et al. 2020) [34]. The shuttle was 65 μm wide, with a planar 11 μm thickness. The T-profile extended 27.5 μm from the back with a width diminishing from 16 μm to 2 μm. This material and profile increased the buckling load of the shuttle by a factor of 13 as compared to a planar silicon shuttle without the T-stiffened profile [34]. This design allowed for array insertion without removal of the tough epineural layer surrounding the DRG, but with presumably less damage to the surrounding tissue. A colorized close-up of the shuttle is shown in Figure 1 (b). The shuttle was glued to the end of the insertion jig prior to adhering the array. The combined array, PCB, jacket, shuttle, and insertion jig will be collectively referred to as the insertion assembly. Figure 1 (c) shows the insertion jig close-up.

### *In Vivo* Deployment

Neural recordings were performed in the DRG of intact, domestic, short-haired adult cats (Liberty Research, Inc., Waverly, NY). All procedures were approved by the University of Michigan Institutional Animal Care and Use Committee, in accordance with the National Institute of Health’s guidelines for the care and use of laboratory animals. Animals were free-range housed prior to use with 0–3 other male cats in a 413 ft^2^ room with controlled temperature (19 °C–21 °C) and relative humidity (35%–60%), food and water available ad lib, a 12 h light/dark cycle, and enrichment via toys and daily staff interaction.

Initial anesthesia was induced with an intramuscular dose of ketamine (6.6 mg kg^−1^)-butorphanol (0.66 mg kg^−1^)-dexmedetomidine (0.033 mg kg^−1^) intramuscular (IM) dose. Animals were intubated, then maintained on isoflurane anesthesia (2%–4%) during the remainder of the procedure. Respiratory rate, heart rate, end-tidal CO_2_, O_2_ perfusion, temperature, and intra-arterial blood pressure were monitored continuously using a Surgivet vitals monitor (Smiths Medical, Dublin, OH). Intravenous (IV) lines were inserted into one or both cephalic veins for infusion of drugs and intravenous fluids (1:1 ratio of lactated Ringers solution and 5% dextrose, 5–30 ml kg^−1^ h^−1^).

A laminectomy (removal of dorsal spinal column bone) was performed to expose the lumbosacral spinal cord and sacral DRG (typically S1–S2). Following laminectomy, the cat’s pelvis was suspended from a custom support frame (80/20 Inc., Columbia City, IN) with stainless steel wire and bilateral bone screws in the superior posterior pelvic crest to minimize spinal motion during breathing and bladder filling. A separate stabilizing frame consisting of optomechanical components (Thor Lab, Newton, New Jersey) and custom 3D-printed components was assembled around the animal to support a 3-axis micromanipulator (502600, World Precision Instruments, Sarasota, FL) and linear actuator (M-235.5DD, Physik Instrumente, Karlsruhe, Germany). The insertion assembly was mounted to the end of the linear actuator, aimed at either the S1 or S2 DRG, and inserted at 2 mm s^−1^ in steps of 5-100 um until fully inserted or beginning to detach from the shuttle. Insertion was monitored with a USB microscope camera. When possible, the array was retracted partially between recording sessions to observe different DRG depths.

### Neural Recording

The reference wire (and ground wire, when not shorted to the reference on the PCB) was connected to a 12-gauge stainless steel needle inserted under the skin on the flank. Neural activity was recorded at 30 kHz using the Ripple Grapevine Neural Interface Processor and associated Trellis software (Ripple Neuro, Salt Lake City, UT). We simultaneously monitored bladder pressure at 1 kHz through the urethral catheter with a pressure transducer (DPT-100, Utah Medical Products, Midvale, UT) and analog amplifier (SYS-TBM4M, World Precision Instruments, Sarasota, FL).

A variety of sensory stimuli were applied to activate sacral afferent neurons, to map the location of different neuronal types along the array and to demonstrate the array’s potential usefulness for neural prosthesis research. To activate skin afferents, the skin was brushed using a cotton applicator in the sacral dermatome associated with the DRG of interest, including regions of the tail, the anus, the perineum, the external meatus of the penis, and the scrotum [36], [37]. These trials typically involved brushing in bouts of 10 s with 10 s rest periods between bouts. Sensory input timing (onset/offset of bouts) was recorded with a push button, which provided a synchronized digital timestamp within the neural recording files. Visceral afferents of the urethra were activated by sliding a catheter back and forth in the orifice. To activate bladder afferents, room-temperature saline was infused through the urethral catheter in sequential boluses of 10 mL each.

For measurements of nerve conduction velocity (CV), electrical stimulation was applied (biphasic, 1:2 charge balanced, cathode-leading, 200 μs pulse-width) at low levels (15–300 μA) to the ipsilateral pudendal nerve via an implanted bipolar nerve cuff (2.0 mm inner diameter Silastic 508-009 tubing; 0.4 mm stainless steel Cooner wire contacts [10]). As an alternative stimulation site, a pair of fine-wire electrodes (stainless steel, 50 μm diameter, Model EMT-2-30, Microprobes, Gaithersburg, MD) was inserted near the anal sphincter and stimulated with a similar waveform at a higher amplitude to generate muscle twitch (0.3-4 mA).

At the end of each experiment, defined as all activities performed in a single animal, euthanasia was achieved with an intravenous dose of sodium pentobarbital (390 mg ml^−1^) under deep isoflurane anesthesia, followed by bilateral pneumothorax. Each experiment, which typically included studies outside of this investigation, lasted approximately 30 hours, of which 4-5 hours was typically allotted for the studies described here. To add context to recordings, DRG from some experiments were removed and fixed in formalin, processed into parafinn blocks, stained with hematoxylin and eosion, and imaged with an inverted microscope (IX83, Olympus, Shinjuku, Tokyo City, Tokyo, Japan), with the brightfield setting at 10 times magnification and Nikon (Minato, Tokyo City, Tokyo, Japan) Element BR 3.2 Software.

### Data Analysis

In order to efficiently handle the large data sets generated by these recordings, we chose to use the open source IronClust suite for MATLAB, which is specifically optimized for high-density probes that oversample individual neurons [38]. Our spike-sorting workflow using IronClust (developed by James Jun and teams at the Janelia Research Campus and the Flatiron Institute) consists of 1) preprocessing, 2) spike detection & feature extraction, 3) density-graph clustering, and 4) manual clustering. We chose IronClust for its real-time processing speed with a GPU and its ability to accurately handle the potential probe drift on flexible probes. IronClust has been evaluated in comparison to several other high-density sorting systems and shown not only to be accurate but to have particular advantages in datasets with probe drift [39]. Each recorded channel was band-pass filtered (300-6000 Hz) and the narrow-band noise peaks were automatically removed in the frequency domain if they exceeded 10 MAD (median absolute deviation) above the median trend curve [40]. Subsequently, the common-mode noise was removed by subtracting each channel by the average across all channels. The remaining motion artifact primarily due to analog-to-digital conversion saturation was rejected by computing the standard deviation of the filtered signals across all channels in each time bin (5 ms duration), and the spike detection was disabled in the time bins exceeding a MAD threshold of 20. The spikes were detected at their negative peaks exceeding 5 MAD threshold [40], and duplicate spikes from the same and neighboring channels were removed if larger peaks were detected within their spatiotemporal neighborhood (50 μm, 0.3 ms). Spike waveforms (1 ms width) surrounding each peak event are extracted from a fixed number of adjacent channels (80 μm). For each spike, we also extracted spike waveforms centered at its secondary peak channel to account for the random jitters of the peak channel due to recording noise and probe drift. Two principal component features were extracted from each channel using a common set of principal vectors for all channels.

In order to handle the probe drift, time bins where the probe occupied similar anatomical locations were grouped together based on the similarity of the 2D histogram of the spike amplitudes and positions. Parameters used were typically those recommended by IronClust developers, either as defaults within the software or in publicly available software evaluations [41]. Anatomical snapshots were computed at regular spike-count intervals such that each snapshot contains an equal number of spike events from all channels (20 s average duration). For each snapshot, a 2D histogram representing the anatomical features was computed by counting spikes based on their amplitude quantiles (8 bins) and center-of-mass positions on the probe. Each time bin was grouped with 14 other time bins exhibiting high similarity scores to form a 300 s average duration. *k*-nearest neighbor (*k*_*NN*_, *k*=30) distances (*d*_*knn*_) were computed between spikes whose peak appeared in channel *c* and time bin *s* with the neighboring spikes whose peak or second peak appeared in channel *c* and time bins that were anatomically grouped with *s*.

Density-graph clustering was performed based on the *k*_*NN*_ [42], [43] by considering a fixed number of local neighbors to achieve a linear scaling. For each spike *j*, the local density score was calculated (*ρ*_*j*_ =1/*d*_*knn,j*_), and the distance separation score (***δ***_*j*_= *d*_*min,j*_ / *d*_*knn,j*_) was calculated where *d*_*min,j*_ is the distance in the principal component feature space to the nearest spike k having a greater density score (min(*dj*_*k*_ | *ρ*_*k*_ >*ρ*_*j*_)). Local density peak points were identified based on a density separation criterion (***δ***>*1*) and the cluster memberships were recursively assigned to the nearest points toward a decreasing density gradient. To minimize false splitting errors due to drift or bursting, units exhibiting similar waveform shapes were merged (Pearson correlation > 0.985). Finally, clusters were manually split, merged, or deleted by using a set of multiple interactive views in the MATLAB-based GUI. Clusters were manually compared based on waveform shape and firing properties across neighboring trials to determine cross-trial repeats to avoid multi-counting individual clusters. Where applicable, we compared cluster centroids by trial type to look for trends in depth (Tukey’s Honest Standard Difference test, α = 0.05). We used a least squares regression to look for trends relating cluster amplitude, channel span, and other features to recording depth.

To understand the relationship between sensory inputs and neural activity, we calculated the correlation of either bladder pressure or cutaneous brushing (coded as a continuous binary on/off signal) with neural firing rate. We used a correlation threshold of >0.2 for bladder pressure or >0.6 for cutaneous brushing to identify related units. The bladder correlation threshold is consistent with previous studies which have utilized correlation thresholds as low as 0.2 for bladder pressure, as even units with low apparent correlation can be useful for decoding pressure from firing rate [2], [10]. The cutaneous correlation threshold is also consistent with or higher than that reported in previous sensory recording studies in DRG [28], [30], [44]. Each correlated cluster was reviewed for visual correlation, and those with only intermittent correlation (correlated during part of a trial but not stable) were excluded. Supplementary Figure 1 shows the effect of changing the correlation threshold on the proportion of clusters identified as bladder or cutaneous units. The signal to noise ratio (SNR) was defined as the unit peak-to-peak amplitude divided by the root mean square voltage of the channel during the entire trial.

For trials with electrical stimulation, a post-stimulus time histogram (PSTH) was generated for each detected unit. If a firing unit had stimulus-locked timing, we used a normal distribution fitted to the PSTH to determine the mean and standard deviation of the delay. To calculate conduction velocity (CV), we assumed a pudendal nerve to sacral DRG length of 9 cm and an anal sphincter to sacral DRG length of 12 cm based on previous measurements [10]. To determine if the CVs of recruited fiber populations differed by stimulation location, we performed a Student’s t-test with α = 0.05.

## Results

### Array Insertion

We attempted insertion of arrays in 6 different feline experiments. The PEG adhesive used to temporarily adhere the array to the shuttle dissolves quickly, and the region around the DRG often had fluid which regularly shifted with breathing. In the first experiment, touching the array to fluid prior to insertion could not be avoided, and the array would not stay adhered for insertion in 2 of 3 insertion attempts. As the temporary adhesion of the array to the shuttle was only briefly successful in experiment 1, and because our primary goal in these experiments was to examine DRG mapping capabilities with the array, we subsequently moved to using cyanoacrylate to bond the array to the shuttle for insertion. In experiment 5, while insertion of the array was achieved, no neural activity was observed. We observed large non-neural artifacts during cutaneous brushing, which suggested that the reference or ground of the array may have been damaged or malfunctioning. In experiment 6, DRG and system movement due to breathing could not be sufficiently eliminated to allow for clean insertion. Impedance of functional electrodes (<1 MΩ; N = 34, 44, 54, 23, 18, and 37 in experiments 1-6 respectively) had a median of 142 kΩ (interquartile range: 364.3 kΩ) when implanted.

### Neural Recording

In this study we recorded high numbers of sensory neurons from feline sacral DRG, identifying single-unit activity from a range of stimuli. While the number of automatically detected units was not quantified, on the order of 100-140 units were identified automatically by IronClust, which was significantly consolidated down to the number of units reported in the Table 1. The span of electrodes with detected units, number of units, peak-to-peak amplitude, and signal-to-noise ratio (SNR) for each experiment are reported in Table 1. The median peak-to-peak amplitude of recorded units was typically on the order of 50-60 μV, though the maximum observed cluster center had an amplitude of 1334 μV (a tonically activated unit in experiment 3 with 2.2 ± 0.48 Hz firing rate, modulated to ~3 Hz by anus brushing). The smallest observed single-unit cluster center with sensory correlation had a mean amplitude of 20.5 μV and SNR of 4.21 (bladder-pressure modulated unit in experiment 2, correlation 0.64). Each cluster was observed on an average of 11.4 ± 5.2 electrodes. The maximum number of sites observed was 26 (in 2 experiments), representing a span of ±500 μm from the center site. This is a larger recording span than we would normally expect from a single firing neural cell body, and the amplitudes of these units were surprisingly not particularly high (57 and 34 μV peak-to-peak; a scrotal brushing and spontaneous unit, respectively). There are a few possible explanations for this. First, our spike sorting software looks for simultaneous deviations on neighboring channels and can therefore detect units that would otherwise be obscured by noise. Additionally, it is known that the path of DRG stem axons can be winding and convoluted [45], so it is possible that near-simultaneous firing detected from stem axon nodes is being picked up by fairly distant channels. Other multi-unit activity was observed with smaller mean amplitude, but the unit shapes were poorly correlated. Bladder pressure related units were observed in 2 of the experiments. An example bladder unit is shown in Figure 2 (a), with the waveforms at the five highest amplitude channels shown on the right. Cutaneous brushing units were observed in all 4 experiments with neural activity. Units were observed with correlation to scrotal brushing, anal brushing, and brushing the dorsal base of the tail. An example unit related to tail brushing is shown in Figure 2 (b).

**TABLE 1:**
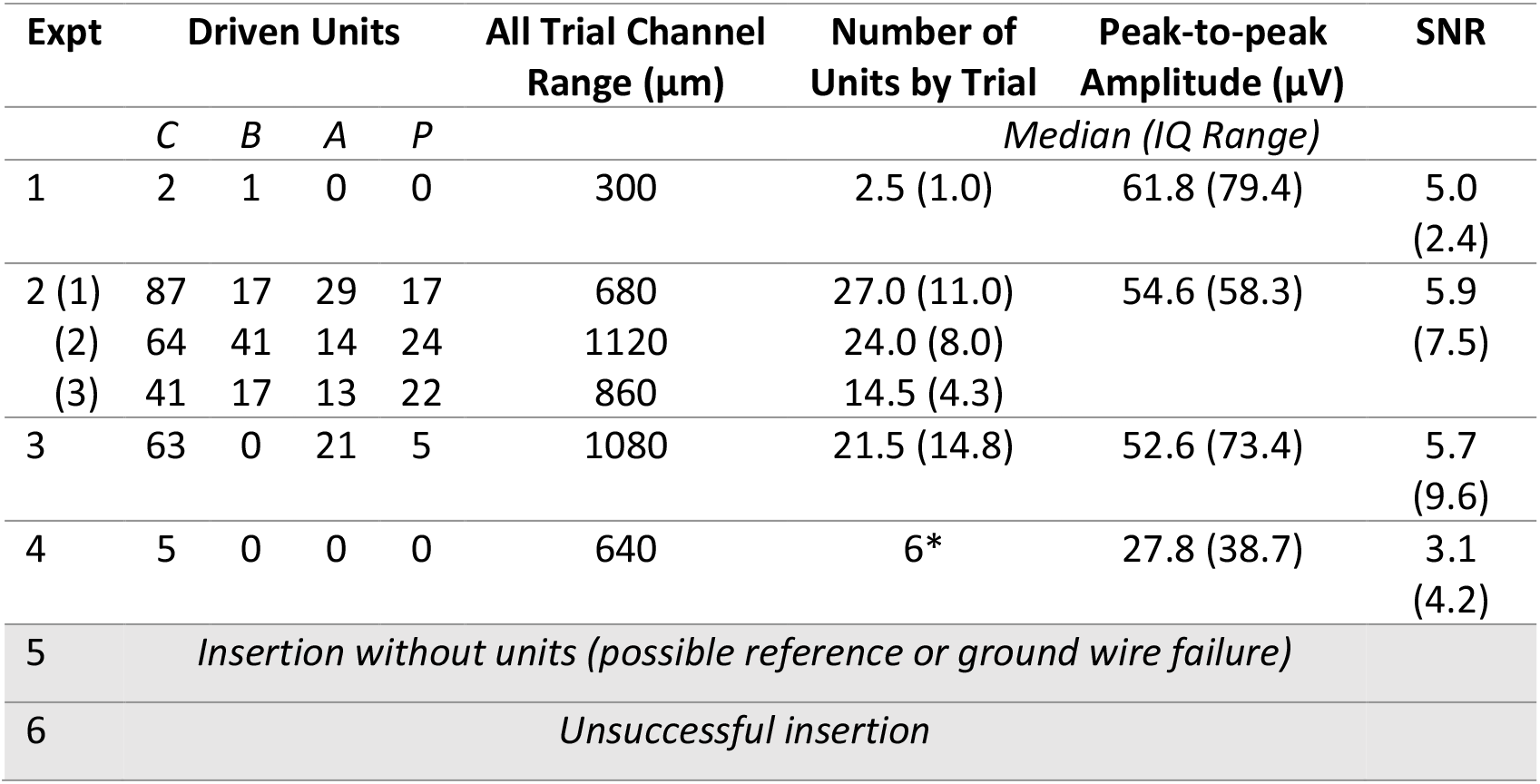
Summary of units recorded during all 6 experiments. Number of driven units is given by unit type: cutaneous (C), bladder (B), electrical stimulation of the anal sphincter (A) or pudendal nerve (P). Number of units by trial, peak-to-peak amplitude, and SNR are given as median with interquartile (IQ) range. For experiment 2, which had three successful positions of a single insertion, select details about each position are given in rows. See Figure 3 for position reference. *In experiment 4, units were only recorded in one trial

**Figure 2:**
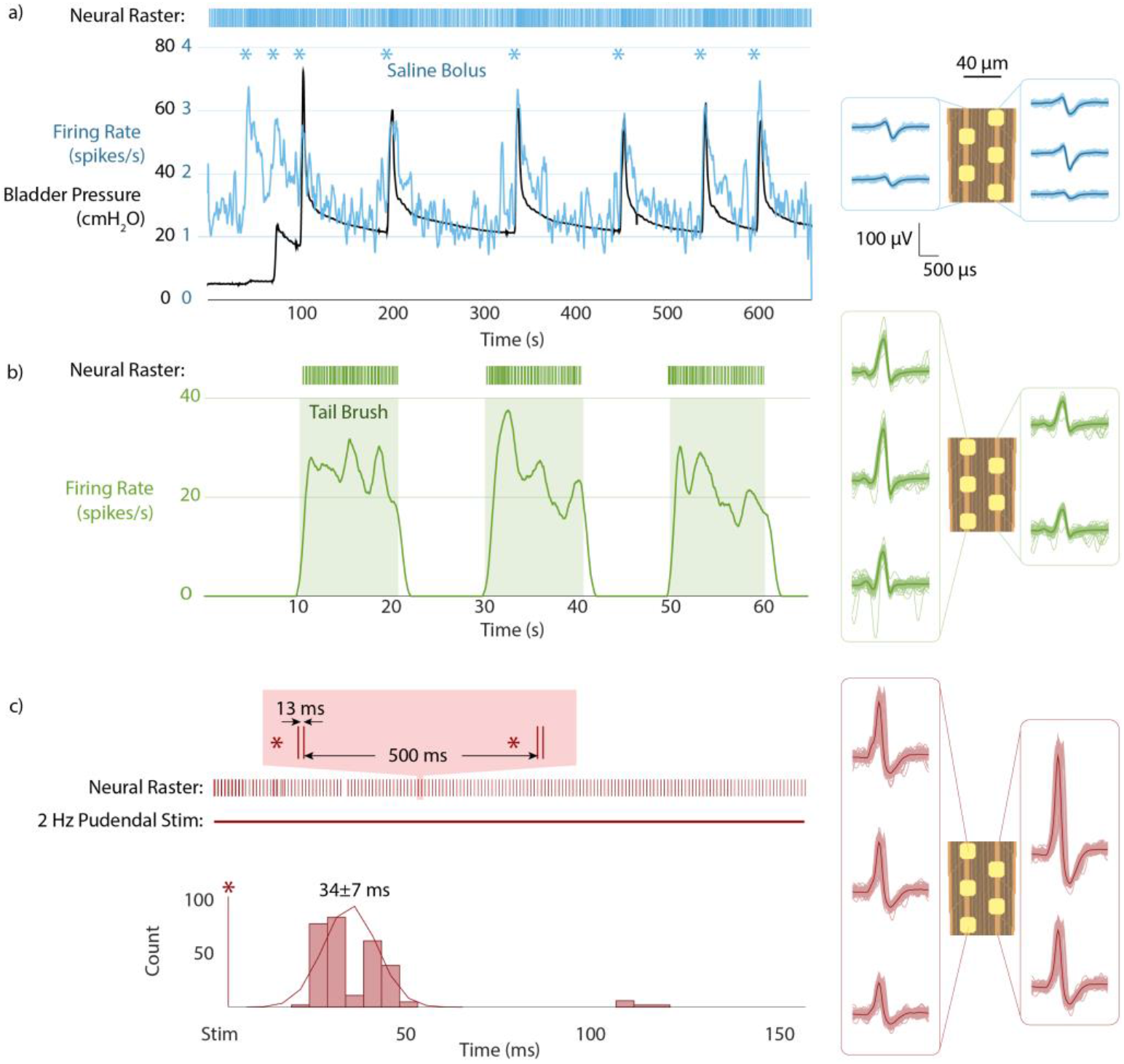
Sample of sensory units recorded from sacral DRG. (a) Bladder-pressure correlated unit from saline bolus fills. Firing rate shown with recorded bladder pressure. Raster plot shows actual spike times. Waveforms shown at right in relation to channels. (b) Tail-brushing correlated unit. (c) Anal sphincter electrical stimulus driven unit (2 Hz, 3.2 mA). Magnified raster plot shows characteristic double-spike response with ~27 then ~41 ms stimulus delay. PSTH shown below neural raster.

Units activated by electrical stimulation of the pudendal nerve or anal sphincter were observed in 2 of the experiments. There was no significant difference in the population of conduction velocities elicited by pudendal or anal stimulation. An example unit is shown in Figure 2 (c), with the associated PSTH showing a delay of 34 ± 7 ms from stimulation to recording. This unit showed a characteristic double spike response to stimulation (anal sphincter, 2 Hz, 3.2 mA), with the first peak around 27 ms and the second around 41 ms. The early peak yields a CV of about 4.4 m/s, which suggests an Aδ-type fiber [46]. There are a number of possible explanations for the second peak. The first peak is most likely a direct activation of the nerve ending by electrical stimulation, and the second likely originates as a result of an ensuing evoked muscle twitch. A variety of single and double-activated units were found in the data set. The longest delay for a directly activated unit was 203 ± 3 ms (CV: 0.44 ± .01 m/s, a pudendal activated C-fiber [46]). The shortest delay for a directly activated unit was 7 ± 0 ms (CV: 12.86 ± 0 m/s, a pudendal activated Aδ-fiber [46]). Other units had a less specific activation tied to stimulation. These units (amplitude on the order of 20-30 μV) were more likely to be active in the period 35-75 ms after a stimulus, but with delay standard deviations of up to 42 ms.

### Recording During Retraction and Breathing

Figure 3 (a) shows the modulated and spontaneous activity recorded at each position of the array in one insertion-retraction sequence during experiment 2. While the exact position of the array relative to the stained section is not known, by comparing the activated regions with the histology we estimated the position of the array in the DRG and the ventral root (VR) below, from which we do not expect to record any sensory-evoked units. Figure 3 (b) shows the putative location of the array relative to the DRG cross section.

**Figure 3:**
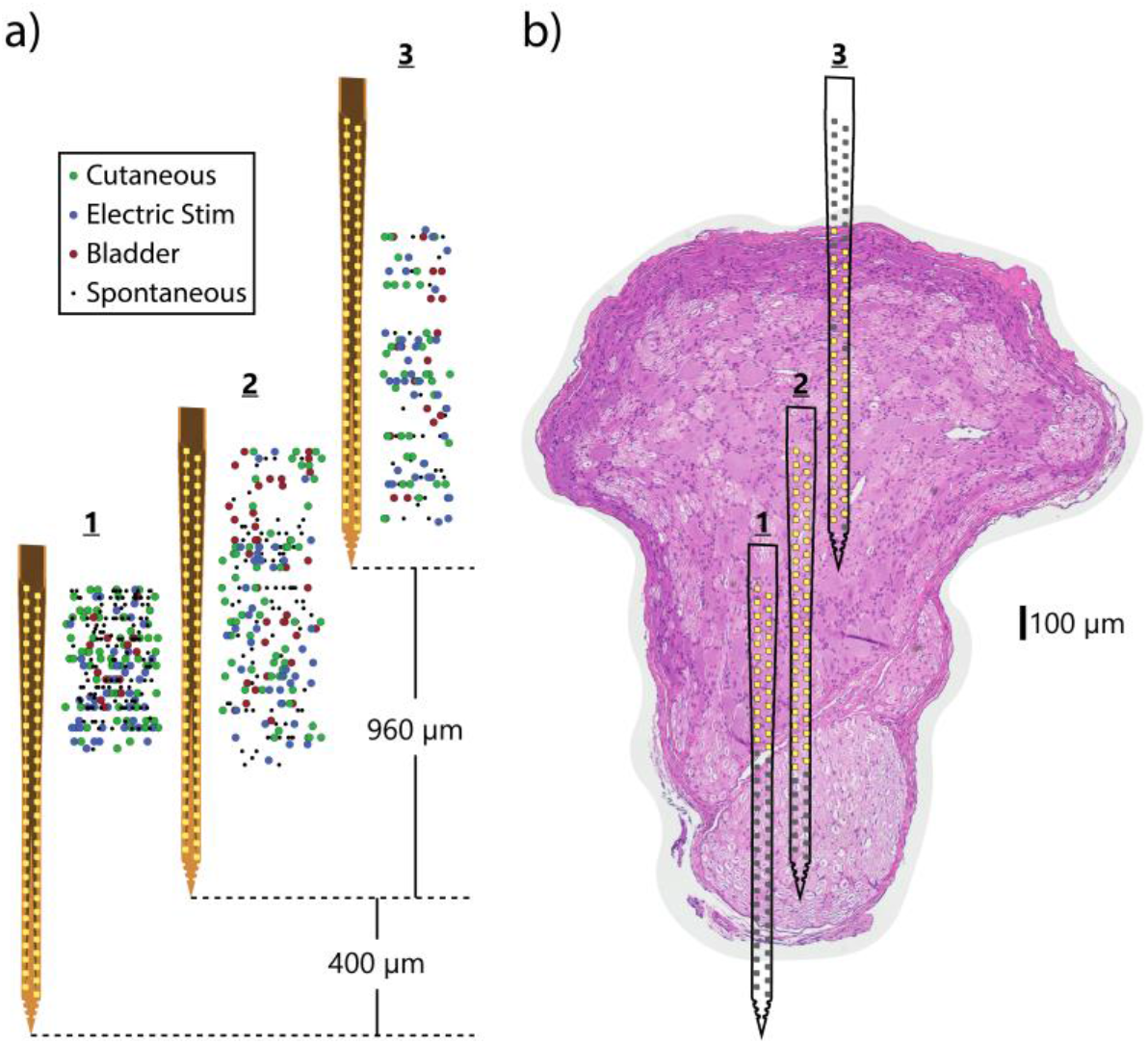
Summary of neural units recorded at different estimated vertical positions in DRG in one experiment. (a) Three different vertical positions of the array in one experiment showing the locations of recorded units (note: horizontal position is not relevant, points jittered for clarity). (b) Putative position of the array relative to histology of sacral DRG from same experiment. Lighter pink region at bottom is ventral root (VR), which does not contain sensory cell bodies for recording. Electrode sites with observed activity are highlighted in yellow, rest are gray. Horizontal position of array does not indicate horizontal movement.

The proportion of cutaneous to overall clusters at positions 1, 2, and 3 were, respectively, 83/286 (29%), 80/269 (30%), and 48/148 (32%). The number and proportion of bladder clusters were 13 (5%), 41 (15%), and 17 (11%). The proportion of electrical stimulation-activated clusters was 46 (16%), 48 (18%), and 49 (33%). The remaining clusters were unrelated to any of the stimulation modalities. Based on the Tukey HSD test, there was not a statistical difference between the average depth of clusters based on stimulation type. The difference between electrical stimulation clusters (1165 ± 490 μm) and bladder clusters (1004 ± 470 μm) was the closest to statistical significance (p = 0.051). The average depth of cutaneous clusters was 1120 ± 465 μm.

Figure 4 (a) shows the number of identified clusters that were detected at each electrode site for the three primary locations in experiment 2. For these array locations, there appeared to be a greater count of clusters closer to the ventral part of the DRG, though this is difficult to evaluate statistically because of the overlap in array positions. Other array positions across our experiments did not have a clear trend. The average waveform peak-to-peak amplitude for each electrode site across these array positions is shown in Figure 4 (b). Visually, there was a greater number of mean waveforms above 100 μV peak-to-peak closer to the ventral part of the DRG in this data, aligning with the greater number of clusters observed in these electrode placements. Least squares regressions of a variety of factors (channels per cluster, waveform amplitude, etc.) did not reveal any trends related to cluster depth in any experiment.

**Figure 4:**
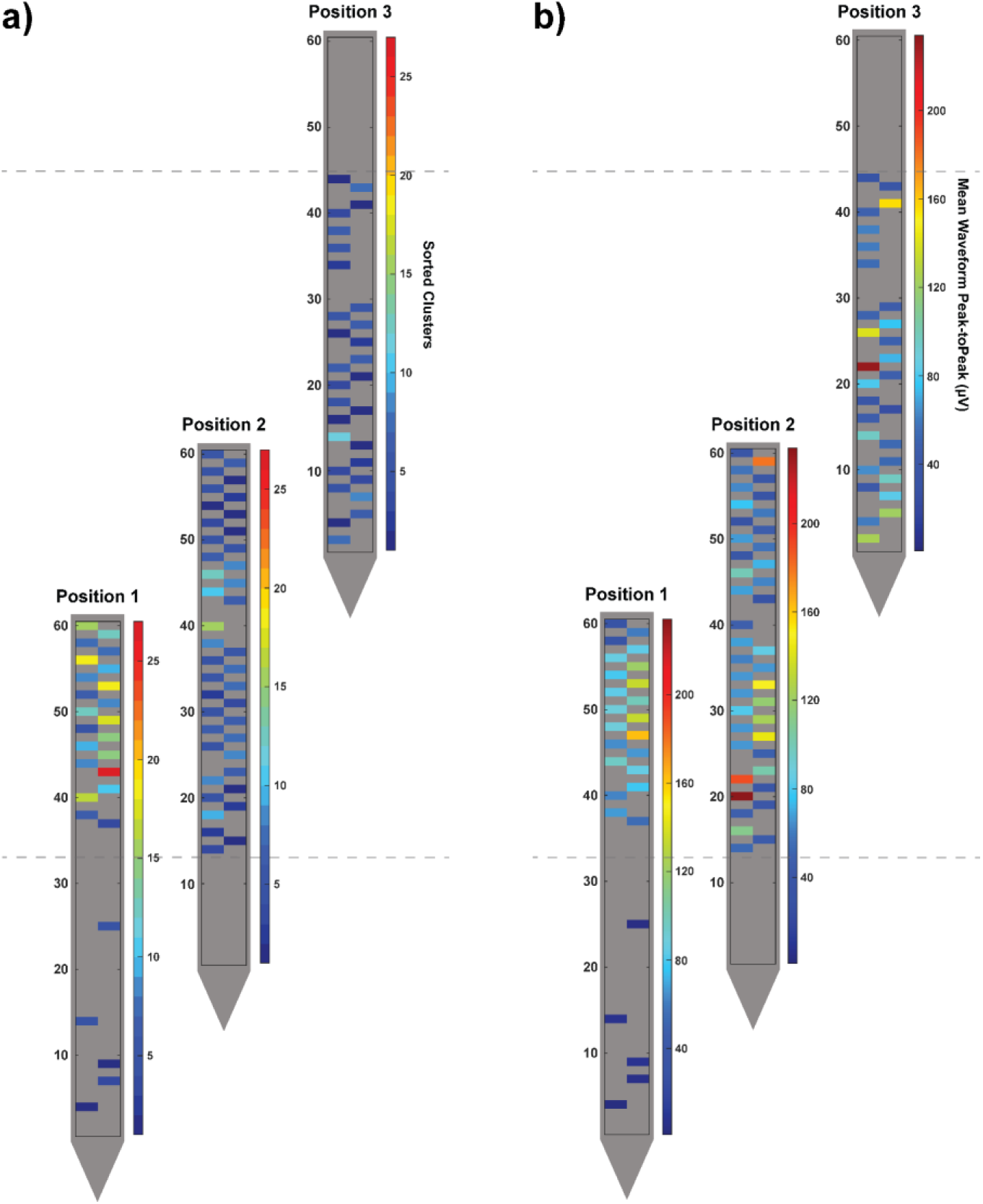
Quantifying observed neural activity across the span of a DRG. a) The number of sorted clusters detected at each electrode site and b) the mean waveform peak-to-peak amplitude at each electrode site are both shown for the three electrode positions in Figure 3. The dotted lines mark the top and bottom of the approximate DRG region where most recordings are expected, per Figure 3.

In the same experiment, we recorded neural activity during 2 mm s^−1^ micromotor retraction of the array between positions, while simultaneously brushing the right side of the scrotum. While noise was too high to discern neural activity during the retraction from position 1 to 2, neural activity was recorded from position 2 to 3. The source of noise during the first retraction but not during the second was not clear. Figure 5 shows putative movement of 8 units throughout the retraction. We observed that the movement of recorded units on the array (~600 μm) does not precisely align with the movement of the array itself calculated from the retraction steps (~960 μm). This is likely a result of slack in the flexible array ribbon. As the array moved relative to neural sources, the detected amplitude and SNR on individual electrodes increases then decreases again. Supplementary Figure 2 shows this expected change in SNR on all channels as the array moves past neural sources.

**Figure 5:**
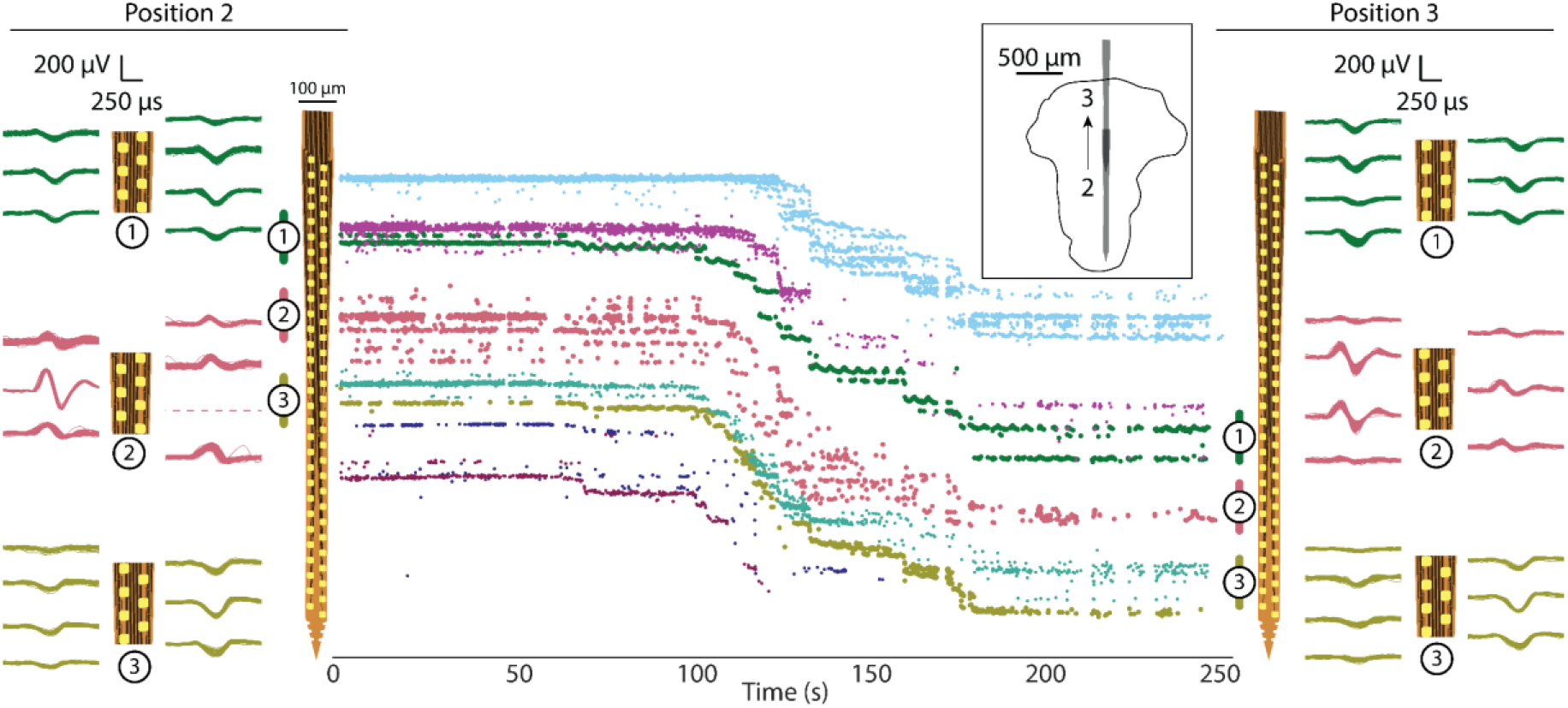
Units tracked across array during withdrawal from position 2 to position 3 (see Figure 3 and inset). As array is withdrawn from DRG, units move relative to the array (toward the tip). Eight units are shown, with movement of approximately 500-600 μm. Waveforms of several of these units are shown at position 2 (left) and position 3 (right) to show similarity of shape.

At certain array positions we observed that clusters would shift among electrode sites in a periodic manner, covering a distance on the array of 10-20 μm. This effect is investigated in Figure 6, for the experimental recording session when the array was retracted in steps (Figure 5). Upon closer inspection we determined that the periodic shifting of the recordings generally cycled at a rate which matched the respiration interval (15 breaths per minute = 1 cycle every 4 seconds). For the blue cluster after 150 s and after 200 s in this sequence, there were non-functioning electrode sites within the vertical span that the cluster covered. This led to gaps within the plotting of cluster locations over time shown within Figure 6. Breathing motion-associated deviation of the other detected clusters were either very weak or non-existent. It is unclear why this difference exists. We speculate that if the electrode array is shifting relative the firing neural body, neural bodies closer to the array will appear to shift more than those further away.

**Figure 6:**
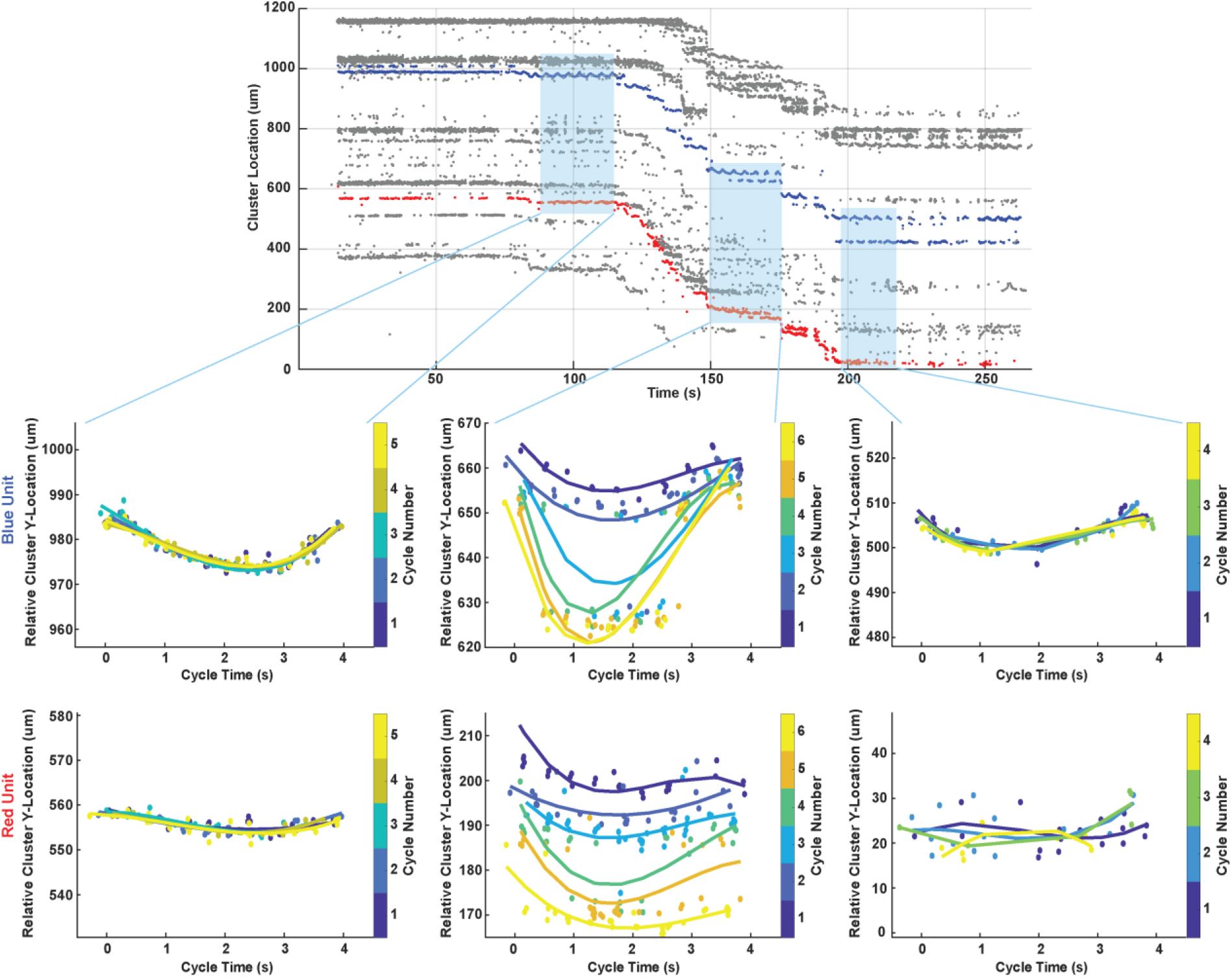
Analysis of breathing effect on clusters shifting among electrode sites, for array movement from position 2 to 3 in Figure 5. Relative vertical locations for two clusters (blue and red in upper plot) are overlaid at four-second intervals per the breathing cycle period for three fixed array locations (light blue shading), showing in some cases a consistent effect on neural recordings. Overlaid on individual data points for each cluster interval cycle is a fit line created with the MATLAB polyfit function. Data points and fit lines are colored per the cycle number key for each sub-plot.

## Discussion

In this study we demonstrated acute high-density recordings from feline sacral DRG. This study is the first to measure neural signals from inside DRG with flexible recording arrays and sets a new milestone for recording density in the peripheral nervous system. Using software specifically designed for sorting high-density neural recordings, we showed that the array was capable of recording neural signals related to bladder pressure and cutaneous brushing in the sacral dermatome, as well as neurons which fire in response to electrical stimulation of the pudendal nerve or anal sphincter. We recorded neural units with peak amplitudes ranging from 20 μV to 1334 μV and tracked neural units while the array was physically moved by utilizing drift-tracking features built in to IronClust. This work shows the potential for high-fidelity interfaces with DRG that can yield new mapping information while being unaffected by relative changes in neuron vertical positions with respect to the array.

A variety of other high-density or special-geometry microelectrodes have been developed for use in the brain [47]. Our work here is an extension of those studies to DRG, examining an over-sampling of local signals to obtain a greater resolution of underlying neural activity. With our high-density probe, we showed that recordings of single units could be achieved on multiple sites for a variety of afferents (Figure 2) and tracked as the array moved over ~1 mm (Figure 5). Units which appeared only on a single channel here would have only a ~15% chance of being recorded with a 400 μm pitch Utah array. One recent technology for high-density recording in the brain is the Neuropixels probe, originally reported by Jun et al. (2017) [48]. This is a stiff silicon electrode array with 960 sites spaced at 20 μm. The array has been demonstrated for high-density single-unit recording in the brains of both head-fixed and chronic freely-behaving mice [48], [49]. While the challenges of implanting, fixing, and recording in DRG are very different from brain, the data processing goals and requirements can be very similar. In fact, the same software suite utilized in our study was also utilized in Jun et al. for faster-than-real-time processing on their very high channel count probes [48].

This study is the also first demonstration of high-density recording at this scale in DRG (40 μm site spacing), and one of very few in the peripheral neural system. Previously, the highest density recordings inside DRG used Utah arrays with site spacing of 400 μm [6], [10], [36], [50], except for a single study with 50 μm-spaced electrodes and no report of unit oversampling [16]. While the types of units recorded from sacral DRG in some of these studies were similar (cutaneous, bladder-related, pudendal-stimulation driven), there was no evidence of unit oversampling on neighboring sites. Higher density recordings have been made from the surface of DRG, down to 25 μm electrode site pitch, [28]–[31], but despite the potential advantages of non-penetrating arrays these were fundamentally limited to recording single-unit activity from the shallowest ~150 μm of the DRG [28]. Slightly higher-density recordings have been reported in the peripheral nervous system. For example, 200 μm pitch Utah arrays have been used to record from the sciatic nerve of rats and the pudendal nerve of cats [51], [52]. Transverse intrafascicular multichannel electrodes (TIME), which penetrate across the nerve axis and could hypothetically be used in DRG, have achieved peripheral nerve recordings with sites spaced at ~230 μm [53]. The recordings achieved in our study therefore set a new milestone for recording density in DRG and the peripheral nervous system. High-density recording in the DRG is not necessary for every application, however. We have previously demonstrated successful neural decoding of bladder pressure using units detected on fewer than 10 channels of a Utah array (in some cases on a third of functional sites), and only one channel may strictly be necessary, if correlation is sufficiently high [2], [3]. Studies implementing closed-loop control of bladder and limb function have also used relatively low-density Utah arrays in DRG during acute, non-survival experiments [3], [5], [44] that would likely lose efficacy in a chronic study. The use of multiple-high density probes inserted across the span of a DRG may yield a richer set of neural signals and, we anticipate, better long-term signals.

This study, in conjunction with a recent report focused on the insertion shuttle technology [34], is the first demonstration of flexible bioelectronics delivered into DRG for neural interfacing. Even in an acute experiment, this approach has potential benefits over the standard Utah array, in potentially reducing bleeding or damage from impact of the pneumatic insertion required for Utah array implant [54]–[56]. In chronic experiments, the mechanical mismatch between stiff materials and soft tissue is expected to result in tissue encapsulation of the shank tip, killing or pushing away neurons in the immediate 40-150 μm vicinity [18], [55], [57]. Our chosen delivery method, the small but stiff diamond shuttle with T-shaped profile, was selected based on the unique challenges of delivering a flexible electrode into DRG through the epineurium. In brain implants, flexible probes can utilize less stiff shuttles because the tough dura mater is typically removed in part prior to insertion for animal models larger than a mouse, exposing the significantly softer parenchyma below [58]. Other studies have addressed delivery of polymer probes by coating them entirely in a variety of soluble stiffening agents, such as PLGA, PEG, or different carbohydrates. All still require removal of the dura [59]. In DRG, however, the tough layer of epineurium cannot be easily removed without damage to the underlying neural tissue. Our UNCD shuttle, with its stiffened T-profile, addresses the need for high stiffness this while maintaining a minimal footprint that reduces damage to underlying tissue and blood vessels, and inserted into both brain and DRG without removal of any outer covering [34]. Shuttles with larger footprints are common, and the use of mechano-adaptive array substrates has also been explored [59].

The primary analysis suite used in this study, Ironclust, is an open-source MATLAB package specifically designed to take advantage of neural unit oversampling to increase the speed and accuracy of spike sorting [38], [39]. While no specific comparison between manual spike sorting was made in this study, a few general observations can be made from the authors’ prior experience with commercial spike sorting software. By considering units identified on clusters of channels, the software removed the effort of separating the same unit on several channels, saving significant time. It also mostly eliminated the danger of yield overestimation. One major benefit for those comfortable with coding (the suite is available in a variety of code languages) was the ability to add features and analysis platforms as needed for a particular study. For example, because of this study’s focus on unit drift, we added a platform to split units not only in principal component space but also based on spatial center. The open-source nature of the project meant that we were able to integrate useful features into the publicly available package which are now available to other researchers.

There was not a consistent clear trend of high cluster count or high signal amplitude near the DRG edge across experiments. Figure 5 suggests a possible increase in detected large-amplitude signals towards the ventral aspect of the DRG within that experiment. Prior work by our group has shown that cell bodies are packed around the DRG perimeter [32], [33], which may yield regions with larger signal amplitude recordings. In our experiments here we may not have been activating all neurons within a region. Additionally, some electrode sites may have been close to active axon nodes, which would cause a lower number of individual clusters to be observed. We were not able to estimate source sizes based on the span of array sites that a cluster appeared on, as our recording array was fixed in a two-dimensions and prevented source localization that we accomplished previously with a flexible array on the curved DRG surface [28]. A better understanding of the types of extracellular waveforms that can be recorded near DRG cell bodies, stem axons, and peripheral axons, through computational modeling, may give more insight into the types of neural elements detected in our recordings.

Loss of signal for chronic intraneural experiments is a common problem [10], [55]. We have observed signal loss or signals reappearing over time with single-site electrode shanks chronically implanted in DRG [10]. This may be due to micromotion of the array, scar tissue development, and/or changing of tissue encapsulation over time. The novel array used in this study could mitigate that effect given its small dimensions and flexibility but needs further evidence. We tracked clusters as the electrode was intentionally pulled rostrally 100s of microns (Figure 5). Furthermore, the high density of our array and the IronClust algorithm allowed for clusters to continually be tracked during small changes in electrode site location. In our case, the relative micromotion of the DRG with respect to the array due to breathing led to visible shifts in cluster locations that were easily tracked across the array (Figure 6). We expect that our combination of a high-density flexible interface with the use of the advanced sorting algorithm for unit tracking will allow for a greater long-term signal yield during chronic *in vivo* studies.

This study, while a successful demonstration of high-density flexible penetrating arrays in DRG, also highlighted some important challenges for future studies, especially any that would proceed to chronic implant and recording. The deployment system was designed to temporarily adhere the array to the shuttle with a dissolvable PEG adhesive during insertion followed by removal of the intact shuttle immediately following array release. Supplemental Video 1 shows a successful deployment of the array *in vivo* to the right S2 DRG in experiment 2. In practice, however, fluid in the surgical cavity could wet the adhesive and cause the array to peel away from the shuttle before insertion. While the system achieved successful delivery in experiment 1, all successful deliveries in following experiments were achieved with permanent cyanoacrylate adhesive to avoid inadvertent early wetting. While we [34] and others [60] demonstrated PEG in rodents, a shuttle inserted through the electrode tip similar to Luan [61] may be more reliable in feline experiments. A future design could use an array with a small loop at the tip to go over the shuttle, which would drive in the array even if the adhesive started to dissolve. The stylet approach has a long history and recently demonstrated on a microscale in the so-called “neural sewing machine” [62].

Another issue with our approach was that, due to large breathing motions following insertion, we not able to successfully withdraw the shuttle without breakage prior to the use of cyanoacrylate. These motions are visible in Supplementary Video 1. We attempted to address this by briefly suspending the breathing cycle during array deployment, but there was insufficient time for the array to fully release from the shuttle before breathing needed to resume. We speculate that this breathing motion and shuttle breakage may account for the array ribbon slack which caused a discrepancy between observed and actual retraction distances (see Results section describing Figure 5). Since the lack of stiff materials in the DRG is one of the primary advantages of our flexible electrodes for chronic use, this issue would need to be solved prior to a long-term implant. One possibility would be to design the shuttle with a controlled breakage point to allow for removal with forceps after the array is securely in place. The natural breakage point of the current shuttle was flush with the DRG surface, making removal difficult. Alternatively, larger “barbs” fabricated as part of the array [63] could hold the array in place during shuttle withdrawal, allowing the shuttle to be removed more quickly.

Assuming these key issues can be addressed, a future chronic study with parallel implant of Utah arrays would be needed to demonstrate the comparative advantage of this technology in both recording longevity and biological response as determined through histological analyses. Continuous neural recording during awake behavior would demonstrate whether the unit tracking demonstrated in this study during array movement would be feasible long-term. This would be a useful feature in developing stable neural decoding for closed-loop neuroprosthesis research. A previous chronic feline study with Utah arrays demonstrated tracking of a bladder DRG neuron over the course of 23 days [10], and computational algorithms can decode bladder pressure from neural firing of one or several units [2], [3], but the long-term stability of these algorithms depends on the ability to monitor multiple bladder neurons over a long period. In terms of biological response, silicon electrode shanks typically produce an active glial scarring region visible on the first day post-implant which stabilizes after 6 weeks, as well as a reduction in neural density within 100 μm of the insertion site [15], [64]. In contrast, thin-film devices have significantly reduced glial scarring even after 4 weeks relative to devices with larger footprints [25], and acute silicon shank stab wounds like those inflicted with our diamond shuttle do not show signs of neural reduction [64]. As previously noted, our shuttle was specifically designed with a tissue-sparing footprint [34]. This said, we did not evaluate the chronic tissue response to our device and insertion method and can only speculate as to these effects in the absence of a chronic study.

We have previously demonstrated with chronically implanted Utah arrays that bladder units in feline DRG are dispersed throughout the entire structure, so there are clear limitations to a single shank technique as presented here [10]. While only single arrays were implanted in the present study, broader mapping of DRG afferents would require multiple arrays implanted in parallel. This could mean a single device with multiple shanks and/or multiple devices implanted next to each other. Further, this penetrating array could be used in conjunction with previously demonstrated surface arrays [28]. To simplify the implant process, it is possible to envision a combined penetrating-surface interface that would unfold onto the DRG surface during insertion. A similar approach has been previously demonstrated for chronic brain recording in a rat model [39]. This approach could provide an anchor for the surface array, a challenge discussed in our previous study [28].

## Conclusions

This study was the first to demonstrate the use of flexible microelectrode arrays to record from within DRG, and to our knowledge the highest-density electrode array reported for use in the DRG or peripheral nervous system. In this study, we recorded a variety of cutaneous, bladder, and electrical-stimulus-driven neural signals from feline sacral DRG in acute anesthetized experiments. We used the high-density data along with specialized open-source software to detect individual neurons recorded on clusters of channels, and to track the “movement” of neural units as an array was slowly withdrawn from the DRG. In the future, we will use these arrays to monitor neurons long-term in awake behavioral studies as we continue to drive the development of neuroprosthetic systems for individuals with neural injury and disease.

## Acknowledgements

We acknowledge all members of Dr. Bruns’ Peripheral Neural Engineering and Urodynamics Laboratory and Dr. Yoon’s Solid-State Electronics Laboratory for valuable assistance in experiments and technology development, especially Aileen Ouyang, Dr. Ahmad Jiman, and Dr. Lauren Zimmerman. Dr. Paras Patel and Elissa Welle assisted in technical aspects of PEDOT coating electrodes. Dr. Mihaly Voroslakos assisted in the design of the array delivery system. Arrays and shuttles were manufactured in the Lurie Nanofabrication Facility at the University of Michigan. Funding for the project was provided by the following: the University of Michigan MiBrain Initiative; the University of Michigan Rackham Predoctoral Fellowship Program; the National Institutes of Health (SPARC awards U18EB021760 and OT2OD023873, and grant 5R21EB020811); the National Science Foundation (Grant # 1653080).

## Author Contributions

Planned study – Z.J.S., K.N., E.Y., J.P.S., T.M.B. Fabricated arrays and shuttles – K.N., J.P.S. Performed surgeries and collected data – Z.J.S., T.M.B. Developed software analysis – Z.J.S., J.J. Analyzed data – Z.J.S., L.R.M., A.S. Drafted manuscript – Z.J.S., L.R.M., T.M.B. Reviewed manuscript and approved final version – Z.J.S., K.N., L.R.M., A.S., E.Y., J.J., J.P.S., T.M.B. Supervised project – E.Y., J.P.S., T.M.B.

## Data Availability Statement

The raw neural and analog recording data that support the findings of this study and will be available with associated metadata from the Blackfynn Discovery platform at doi: 10.26275/vzxw-kwdu after curation by the NIH SPARC Data Resource Center. This includes all data used to generate Figures 2-6, as well as summary data.

## Code Availability Statement

All analysis in this study was performed using ironclust, a neural spike sorting package for MATLAB developed at Flatiron Institute, based on JRCLUST (Janelia Rocket Cluster). The current version of this software, which includes features developed for this study, can be found at https://github.com/flatironinstitute/ironclust.

## Competing Interests

T.M.B. is a named inventor on a granted patent (US9622671B2; assigned to University of Pittsburgh) which is on the monitoring of physiological states via microelectrodes at DRG. The authors declare no other personal or institutional interest with regards to the authorship and/or publication of this manuscript.

## Notes

https://doi.org/10.26275/vzxw-kwdu

### Competing Interest Statement

T.M.B. is a named inventors on a granted patent (US9622671B2; assigned to University of Pittsburgh) which is on the monitoring of physiological states via microelectrodes at DRG. The authors declare no other personal or institutional interest with regards to the authorship and/or publication of this manuscript.

### Summary of Updates

Revision following peer review comments.

