## Supplementary Figure for "High-density Neural Recordings from Feline Sacral Dorsal Root Ganglia with Thin-film Array"

Supplemental Video 1: Insertion of array *in vivo* into right S2 DRG of experiment 2. Video is labeled to indicate location and orientation of spinal cord and bilateral S2 DRG. Visible at right is an array mounted to a nanocrystalline diamond shuttle at the end of the 3D printed insertion jig. Breathing movement is visible within the first 10 seconds of the video. Shuttle is advanced stepwise toward the DRG at 17 seconds and enters the DRG at 41 seconds. This array insertion corresponds to Position 1 in Figures 3-6.

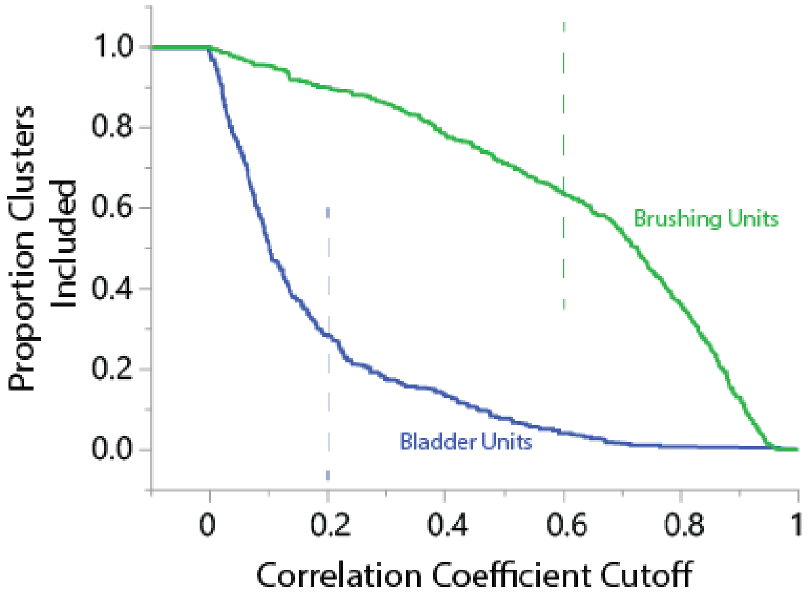


Supplemental Figure 1: Illustration of the effect of changing the chosen correlation coefficient (CC) on the proportion of clusters included as “bladder” or “brushing.” Plot was constructed by calculating absolute value of CC between each cluster detected in either trial type, then counting the proportion of those units clusters with CC below the CC cutoff. For reference, a total of 495 clusters were detected in brushing trials, and a total of 302 clusters were detected in bladder trials. The vertical dashed lines intersecting each curve represent the cutoff value reported in the Results.

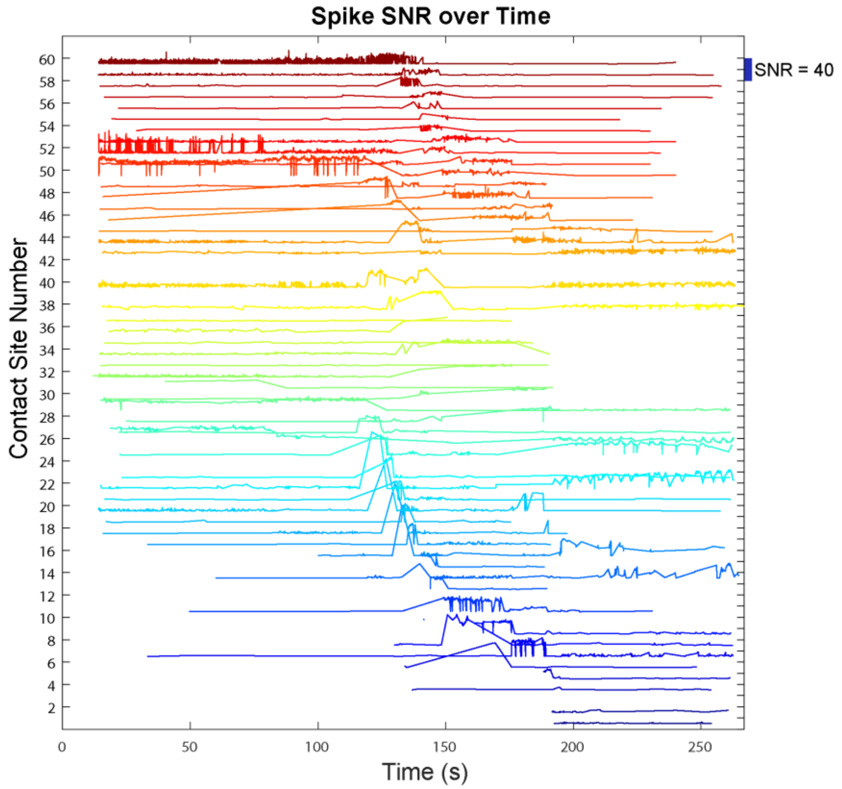


Supplemental Figure 2: Line plot showing the SNR for each spike detected by each contact site over time for the trial shown in Figure 4. Scale bar on the top right shows magnitude of SNR. Generally, SNR increases then decreases as the electrode contact moves past a cluster center.
